# Computationally Reconstructing Cotranscriptional RNA Folding Pathways from Experimental Data Reveals Rearrangement of Non-Native Folding Intermediates

**DOI:** 10.1101/379222

**Authors:** Angela M Yu, Paul M. Gasper, Eric J. Strobel, Kyle E. Watters, Alan A. Chen, Julius B. Lucks

## Abstract

The series of RNA folding events that occur during transcription, or a cotranscriptional folding pathway, can critically influence the functional roles of RNA in the cell. Here we present a method, Reconstructing RNA Dynamics from Data (R2D2), to uncover details of cotranscriptional folding pathways by predicting RNA secondary and tertiary structures from cotranscriptional SHAPE-Seq data. We applied R2D2 to the folding of the *Escherichia coli* Signal Recognition Particle (SRP) RNA sequence and show that this sequence undergoes folding through non-native intermediate structures that require significant structural rearrangement before reaching the functional native structure. Secondary structure folding pathway predictions and all-atom molecular dynamics simulations of folding intermediates suggest that this rearrangement can proceed through a toehold mediated strand displacement mechanism, which can be disrupted and rescued with point mutations. These results demonstrate that even RNAs with simple functional folds can undergo complex folding processes during synthesis, and that small variations in their sequence can drastically affect their cotranscriptional folding pathways.

**Highlights:**

- Computational methods predict RNA structures from cotranscriptional SHAPE-Seq data
- The *E. coli* SRP RNA folds into non-native structural intermediates cotranscriptionally
- These structures rearrange dynamically to form an extended functional fold
- Point mutations can disrupt and rescue cotranscriptional RNA folding pathways

## Introduction

It has long been appreciated that the diverse functions of RNAs are due to specific structures that mediate interactions within the cell (Cech and Steitz, 2014; Sharp, 2009; Strobel et al., 2016). RNA structures begin to form during transcription, where nascent RNA molecules exiting RNA polymerase (RNAP) can transition through intermediate structural states that can ultimately influence and determine RNA function (Heilman-Miller and Woodson, 2003; Kramer and Mills, 1981; Pan et al., 1999; Wong et al., 2007). Because RNA folding generally occurs faster than transcription (Mustoe et al., 2014; Roberts et al., 2008), the 5’ to 3’ directionality of RNA synthesis guides an order of folding, or cotranscriptional ‘folding pathway’ (Pan and Sosnick, 2006). Cotranscriptional folding occurs every time an RNA is transcribed, and can be critically important for essential catalytic RNAs to fold correctly, for riboswitches to make regulatory decisions, for coordinating macromolecular assembly of the ribosome, and to facilitate mRNA processing (Al-Hashimi and Walter, 2008; Fedor and Williamson, 2005; Meyer et al., 2011; Pan and Sosnick, 2006; Russell et al., 2002; Saldi et al., 2018; Serganov and Nudler, 2013; Wickiser et al., 2005).

Despite its importance, we still lack a complete understanding of the dynamic, non-equilibrium folding pathways that RNAs undergo during transcription. Pioneering studies showed that polymerase speed and the order of the RNA sequence elements are important for establishing functional folds of RNA enzymes (Heilman-Miller and Woodson, 2003; Pan et al., 1999). Later, enzymatic RNA structure probing and oligonucleotide hybridization was used to generate structural models of cotranscriptional folding processes (Wong et al., 2007). Single-molecule force measurements have also been used to track major folding events of regulatory riboswitches in real time (Frieda and Block, 2012). In order to complement these approaches with higher resolution structural information, we recently developed cotranscriptional SHAPE-Seq that captures nucleotide-resolution flexibility data for each length of a nascent RNA in stalled transcription elongation complexes using high throughput RNA chemical probing (Watters et al., 2016a). While these experimental methods are becoming increasingly powerful, the data generated is complex and cannot directly obtain specific RNA structure models from the data alone.

Computational RNA folding algorithms are important tools for generating models of RNA structure and folding. Some of these algorithms, such as CoFold (Proctor and Meyer, 2013), modify minimum free energy (MFE) folding calculations to capture some cotranscriptional folding effects. Kinwalker (Geis et al., 2008) and others such as KineFold (Xayaphoummine et al., 2005), RNAKinetics (Danilova et al., 2006), and BarMap (Hofacker et al., 2010) use stochastic simulations of RNA folding with growing chain length to model cotranscriptional folding pathways. Comparative methods have also been developed such as TRANSAT which utilizes multiple sequence alignments and evolutionary trees to capture potentially conserved transient structures (Wiebe and Meyer, 2010; Zhu et al., 2013). These algorithms have been applied to several important biological phenomena, such as kinetic trapping in the Hepatitis δ ribozyme (Isambert and Siggia, 2000), regulation by the adenine-sensing riboswitch (Geis et al., 2008), and folding pathways of the *E. coli* leader sequence of tRNA^phe^ (Hofacker et al., 2010). In some cases, these algorithms have even been used to design new RNAs to fold into different structures depending on sequence orientation (Xayaphoummine et al., 2005), and to design synthetic RNA regulatory switches that regulate transcription *in vitro* (Alexandre et al., 2009; Wachsmuth et al., 2013)

While *in silico* cotranscriptional folding predictors show great promise, the algorithms could benefit from high resolution experimental studies to corroborate, guide and improve their predictions. It is well established that RNA chemical probing data can be used as restraints in computational RNA folding algorithms to improve consistency between equilibrium predictions and experimental measurements (Deigan et al., 2009; Hajdin et al., 2013; Loughrey et al., 2014; Washietl et al., 2012; Watters et al., 2016b). The existing RNA secondary structure prediction algorithms that can use experimental probing data as input can be placed into two groups that either: (1) directly modify the free energy model used in the folding calculation (Deigan et al., 2009; Lorenz et al., 2016a; Lorenz et al., 2016b; Lorenz et al., 2016c; Washietl et al., 2012; Wu et al., 2015) or (2) select a structure from a set of possible structures generated by unmodified folding calculations (Ding et al., 2004; Ouyang et al., 2013; Tan et al., 2017). These methods have shown great improvements in structure prediction, with prediction accuracies increasing by two-fold in some cases (Deigan et al., 2009). Despite this success, these algorithms were developed to model RNA folding in equilibrium conditions, and efforts to predict cotranscriptional folding pathways from chemical probing data have only recently begun (Li and Aviran, 2018).

Here we develop a method called Reconstructing RNA Dynamics from Data (R2D2). R2D2 uses nucleotide-resolution chemical probing data as inputs to reconstruct models of secondary and tertiary RNA cotranscriptional folding pathways. Secondary structure reconstruction for each length of a nascent RNA begins by sampling possible structures using RNA sequence information and cotranscriptional SHAPE-Seq data, and then selecting sampled structures that are most consistent with the experimental data using an optimized distance metric. This secondary structure reconstruction is then used as a starting point for all-atom molecular dynamics simulations to generate three dimensional models of cotranscriptional folding transitions observed between specific predicted intermediate states.

We applied R2D2 to the *Escherichia coli* Signal Recognition Particle (SRP) RNA, a highly conserved non-coding RNA that is also known as 4.5S RNA and is found in all kingdoms of life (Rosenblad et al., 2009). The SRP RNA binds to protein cofactors to form the Signal Recognition Particle, which recognizes nascent signal peptide sequences and delivers ribosome-nascent chain complexes to a membrane for translocation through docking to the SRP receptor. SRP RNA thus plays a critical role in protein biogenesis in the cell.

The *E. coli* SRP RNA is a powerful model for studies of nascent RNA folding because previous studies indicate that it rearranges from a non-native intermediate hairpin fold into an extended helical structure (Wong et al., 2007) that resembles the native functional structure (Batey et al., 2000) during its transcription. We therefore sought to apply R2D2 to reconstruct secondary structure folding pathway models of this model system to uncover mechanistic insights into this rearrangement process. Our secondary structure predictions informed the identification of a single nucleotide mutation within the non-native intermediate hairpin that disrupts the cotranscriptional rearrangement of the SRP RNA. We then performed all-atom molecular dynamics (MD) simulations to assess possible mechanisms for the native sequence rearrangement and insights into how a single mutation can disrupt this process. The simulations suggest that the rearrangement can proceed via a toe-hold mediated strand displacement mechanism, which requires flexibility in the non-native intermediate hairpin that is abolished by the point mutation. We also created point mutations to re-introduce flexibility into the non-native intermediate hairpin which rescued the ability of the SRP RNA to cotranscriptionally rearrange into its native fold. While this work was being performed, several of our structural predictions were corroborated by an independent study which used a high-resolution optical tweezers instrument to follow in real-time and on a single-molecule scale the cotranscriptional folding of the same SRP RNA sequences (Fukuda et al., Submitted). Overall this demonstrates the sensitivity of the cotranscriptional folding pathway of a highly conserved and essential non-coding RNA to small sequence changes, and efficient ways by which RNAs may rearrange out of non-native intermediate folds during transcription.

## Results

### A sample-and-select approach to reconstructing RNA folding pathways from experimental data

We developed a method to merge computational RNA secondary structure algorithms with nucleotide-resolution experimental datasets generated from cotranscriptional SHAPE-Seq that probe nascent RNA structure (Figure 1). Cotranscriptional SHAPE-Seq begins with *in vitro* transcription of a DNA template library that directs the synthesis of each intermediate length of a target RNA using RNA polymerase (RNAP) roadblocks (Strobel et al., 2017; Watters et al., 2016a). Transcription from this template library generates nascent RNAs of all intermediate lengths of the target sequence, which are rapidly probed with the fast-acting SHAPE reagent benzoyl cyanide (BzCN) (reaction t_1/2_ of 250 ms) to covalently modify the RNA according to its structure (Mortimer and Weeks, 2007). Library preparation, sequencing and bioinformatics analysis is then used to generate a SHAPE-Seq reactivity spectrum for each intermediate length of the nascent RNA (Watters et al., 2016a). RNA nucleotides that are unconstrained by secondary or tertiary structure are more susceptible, or reactive, to modification (Aviran et al., 2011; Bindewald et al., 2011; Deng et al., 2016; Sukosd et al., 2013). Cotranscriptional SHAPE-Seq thus generates chemical probing data for each nucleotide of each nascent intermediate length RNA species (Figure 1A).

**Figure 1.**
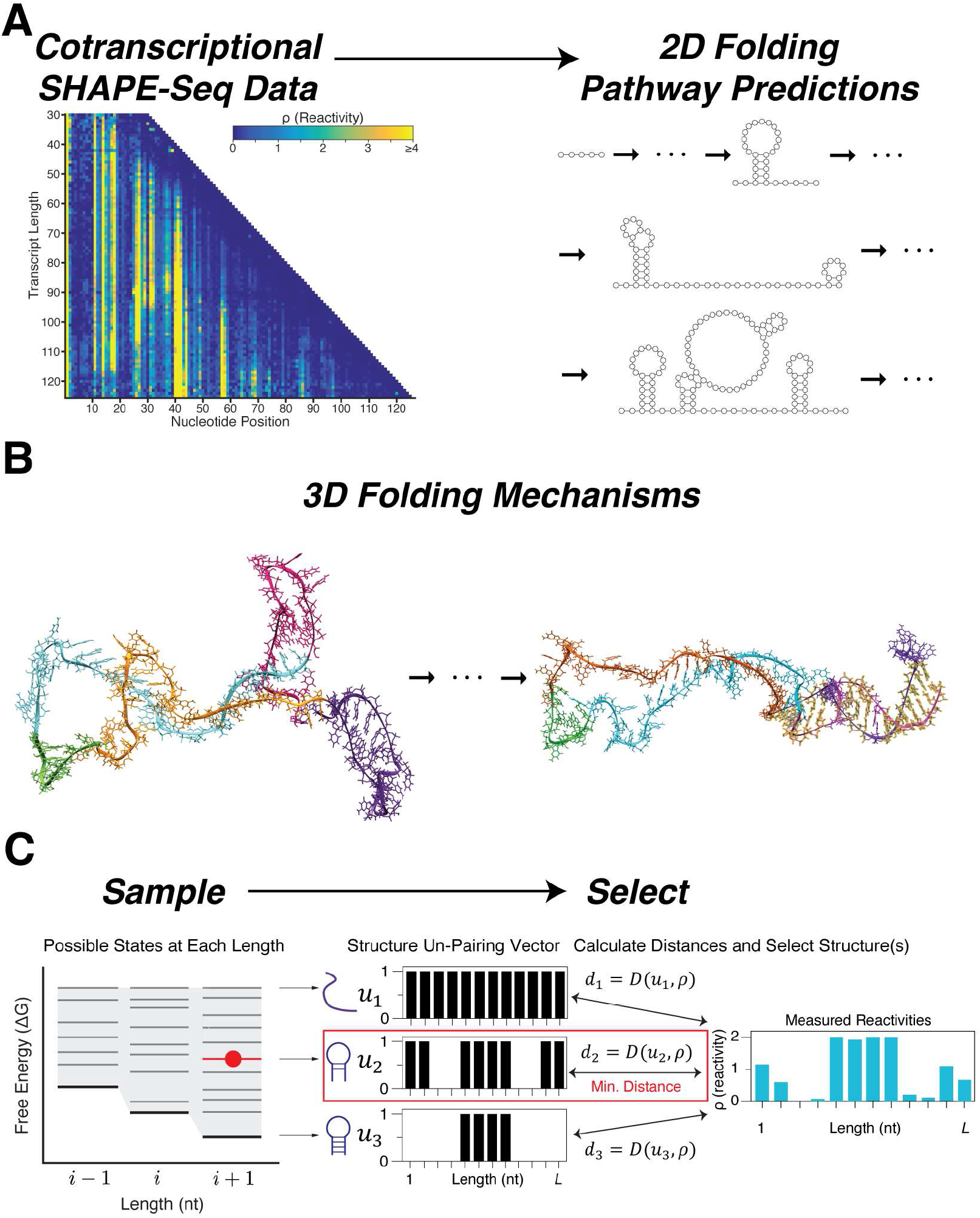
Overview of the Reconstructing RNA Dynamics from Data (R2D2) approach. **(A)** Schematic of the secondary structure prediction method in R2D2. First, cotranscriptional SHAPE-Seq is used to determine reactivities at each nascent transcript length. These reactivities are used to generate 2D (secondary) structures along these transcript lengths. **(B)** 3D simulations are then used to determine the feasibility of structural transitions between specific states within the 2D predictions. **(C)** Outline of the secondary structure prediction method. First, potential RNA structures are statistically sampled for every nascent RNA length. For each length, structures are tested for consistency with the reactivity data at that length, and the most consistent structure is selected. The collection of structures over all of the nascent RNA lengths are then used as potential structures found in the RNA’s cotranscriptional folding pathway.

Inspired by SeqFold and Sfold (Ding et al., 2004; Ouyang et al., 2013), we chose to use a sample-and-select method to reconstruct secondary structure folding intermediates within R2D2. The R2D2 sample-and-select method consists of two steps: (1) generate a set of possible structures at each intermediate nascent RNA length by *sampling* candidate structures from the sequences alone; and (2) computationally *select* the most likely structure(s) using observed experimental data (Figure 1C). Comparison between SHAPE-Seq reactivities and sampled structures with a ‘distance’ metric that reflects how similar reactivity patterns are to candidate secondary structures is used as our method to *select* structures that are most consistent with the data at each nascent RNA length (Figure 1C, Table S1).

To generate candidate structures, the *sample* method statistically samples structures with a large sample size using the *partition* and *stochastic* functions of the RNAstructure suite of computational secondary structure prediction tools (Harmanci et al., 2009; Mathews, 2004; Reuter and Mathews, 2010). The *partition* method takes as an input the RNA sequence and folding parameters, and uses them to calculate the secondary structure partition function for that sequence. The partition function for an RNA sequence describes how a population of RNA molecules with that sequence partitions into an ensemble of different structures in equilibrium, with each structure occurring with a Boltzmann probability. The *stochastic* method then uses this partition function to stochastically generate RNA structures according to their equilibrium Boltzmann probabilities – i.e. lower free energy structures are generated more frequently than higher free energy structures. Thus repeated application of the *stochastic* method can generate a set of possible candidate structures the RNA molecule may exist in during the cotranscriptional SHAPE-Seq experiment. In order to increase the diversity of structures sampled, we applied three variations of the *partition* method that incorporated experimental SHAPE reactivities in different ways to sample 150,000 structures for each length (Methods).

In order to select structures from this sampled set, we developed six metrics to calculate a distance between a given SHAPE-Seq reactivity spectrum and a given RNA secondary structure (Table S1, Methods). Structures were *selected* from a candidate sampled set by minimizing the distances calculated between the reactivity spectrum and the candidate structure set (Figure 1C). By applying this selection at each nascent RNA length, we could reconstruct possible folding intermediates that were most consistent with the experimental data.

### Benchmarking sample-and-select on equilibrium refolding data

We next assessed the accuracy of each proposed distance metric. As there are currently no established benchmarks for cotranscriptional folding predictions, we instead assessed distance metrics by predicting the equilibrium folds of an established benchmark panel of RNAs using SHAPE-Seq equilibrium refolding data (Deigan et al., 2009; Hajdin et al., 2013; Loughrey et al., 2014). Each distance metric contains several parameter values that are used to determine how the SHAPE reactivities are compared to sampled RNA structures: *ρ_max_* and *ρ_c_* determine cutoffs in reactivity values and *α* weighs the contributions from paired vs. unpaired positions (Methods). For each of the six distance functions, we determined the optimal values of the three fit parameters by applying the sample-and-select method on a panel of RNAs previously used to benchmark equilibrium SHAPE-directed secondary structure prediction algorithms: 5S rRNA (*E. coli*), adenine riboswitch (*V. vulnificus*), P4-P6 tetrahymena group I intron ribozyme, TPP riboswitch (*E. coli*), cyclic di-GMP riboswitch (*V. cholera*), and tRNA^phe^ (*E. coli*) (Deigan et al., 2009; Hajdin et al., 2013; Loughrey et al., 2014) (Table S1). The best performing parameter sets performed comparably to RNAstructure-Fold, a widely used RNA secondary structure prediction algorithm, when given SHAPE data (Table S2, Methods).

### *Reconstructing the secondary structure cotranscriptional folding pathway of the* E. coli *signal recognition particle RNA sequence*

We next applied the R2D2 sample-and-select method to our previously published cotranscriptional SHAPE-Seq dataset for the SRP RNA sequence (Watters et al., 2016a). For these experiments, the SRP RNA sequence was obtained from a previous investigation on its folding pathway (Wong et al., 2007), which replaced the 24nt leader sequence that is cleaved off during SRP RNA precursor processing in the cell (Li and Altman, 2003), with three nucleotides (AUC) on the 5’ end of the sequence. We therefore used this modified SRP RNA sequence in this study. In order to apply R2D2 sample-and-select to this dataset, we removed the last 14nts from each 3’ end of the RNA sequence in cotranscriptionally probed datasets to account for the RNA polymerase footprint (Komissarova and Kashlev, 1998; Watters et al., 2016a). To compare cotranscriptional predictions to those made from equilibrium refolded datasets that do not contain a polymerase footprint, we shifted the cotranscriptional transcript lengths by 14nt to compare equal lengths of the RNA sequence that is free to fold from each experimental dataset. To visualize R2D2 predictions at each nascent RNA length, we visualized the selected structures by plotting their free energies (ΔG) and connected all possible paths between selected structures for visual convenience, noting that connections do not imply transition probabilities between states (Figure 2A). Notably, we observed that distinct structures can have the same minimum distance to the experimental data, which may reflect a mixed population of RNA states at specific lengths (Li and Aviran, 2018; Spasic et al., 2017; Watters et al., 2016a). We therefore chose to leave these multiple structures as possibilities that are equally consistent in some way with the data.

**Figure 2.**
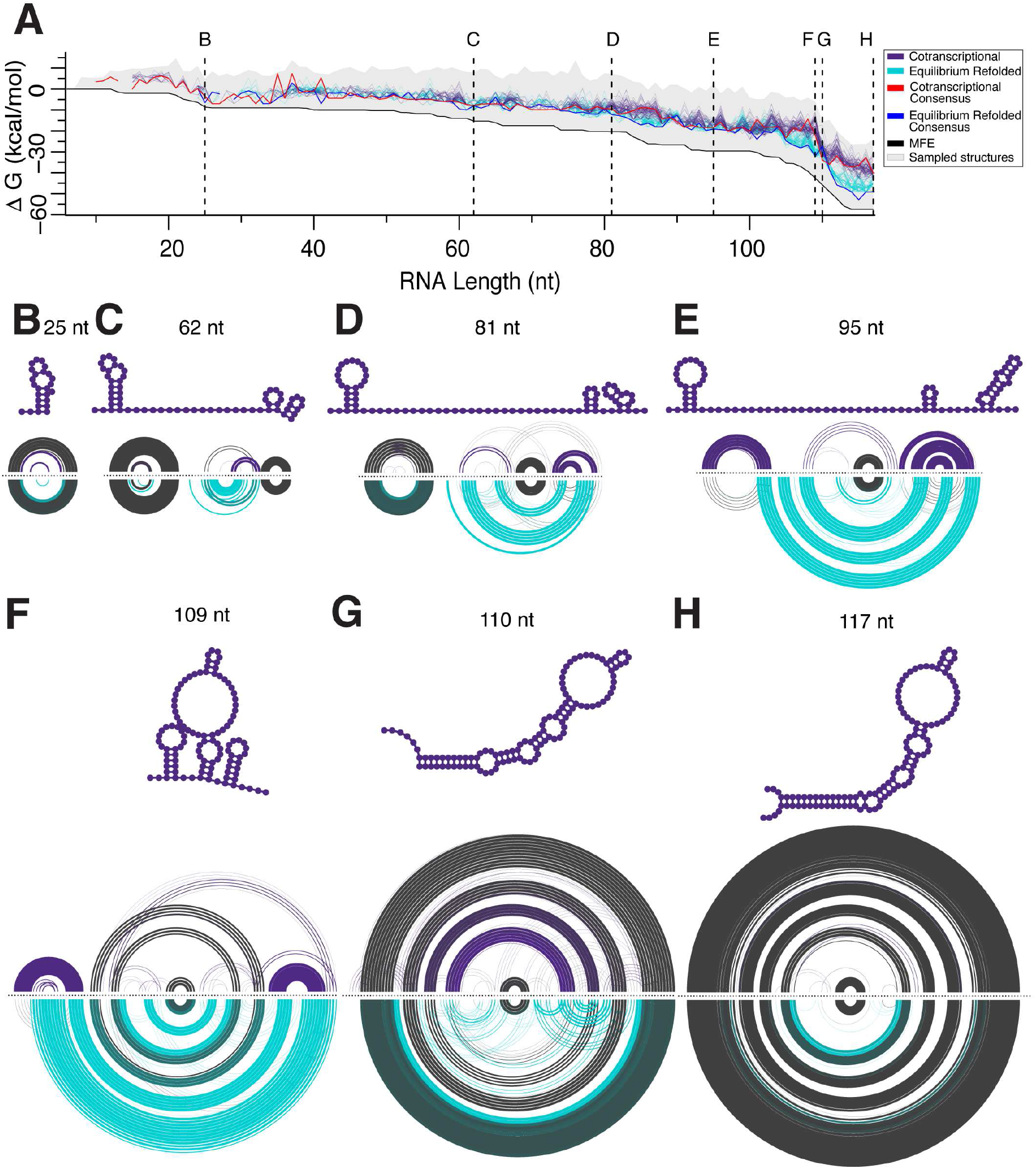
R2D2 2D pathway predictions for the *E. coli* SRP RNA sequence. **(A)** Secondary structure predictions by R2D2 on cotranscriptional and equilibrium refolded SHAPE-Seq data of the *E. coli* SRP RNA sequence. For each dataset, 100 folding pathway predictions were performed and plotted according to the free energy (ΔG) of the RNA structures predicted along the cotranscriptional (purple) or equilibrium refolded (turquoise) pathway. The range of ΔG’s sampled at a given length are represented by grey shading, while the specific ΔG of the structures chosen at each length are represented by dots. For visual convenience, dots are connected by lines to view possible free energy changes along the folding trajectory. Consensus structure lines connect free energies of structures containing base pairs that occur in over 50% of the 100 iterations of R2D2 performed on the cotranscriptional (red) and equilibrium-refolded (blue) SHAPE-seq data. The black line connects the minimum free energy structures in the sampled set. Seven lengths of 2D R2D2 predictions are highlighted: **(B)** 25 nt, **(C)** 62 nt, **(D)** 81 nt, **(E)** 95 nt, **(F)** 109 nt, **(G)** 110 nt, **(H)** and 117 nt. For each length, the 100 selected structures are represented as RNAbow plots where base pairs are drawn as arcs with the width of the arc indicating higher prevalence of the base pair. Colored arcs show base pairs that are more frequent in either cotranscriptional (purple) or equilibrium (turquoise) predictions, while grey arcs show base pairs that are similar between the two sets of predictions. The consensus structures from cotranscriptional predictions are shown above each RNAbow plot. We shifted the cotranscriptional transcript lengths by 14nt to compare equal lengths of the RNA sequence that is free to fold from each experimental dataset. Data plotted in this figure is from cotranscriptional SHAPE-Seq replicate 1.

Despite diversifying our sampling procedure 150-fold over some previous sample-and-select methods (Ding et al., 2004; Ouyang et al., 2013), it is intractable to generate an exhaustively complete set of candidate structures at each length due to the slow convergence of the stochastic sampling method (Figure S1). Thus, repeated applications of sample-and-select may generate different unique sets of candidate structures that can be consistent with the data. To incorporate this variability in sampling, we ran 100 iterations of R2D2 sample-and-select on each SHAPE-Seq dataset to give a family of possible intermediate folding states (Figure 2A). We then applied this method to cotranscriptional SHAPE-Seq datasets of the SRP RNA sequence, as well as SHAPE-Seq datasets from experiments performed on an equilibrium refolded population of the same SRP RNA sequence intermediates to compare out-of-equilibrium to equilibrium predictions of intermediate states (Figure 2, Figure S2). Overall, we see that cotranscriptional and equilibrium predictions are similar for short RNA lengths, but diverge as the RNA length increases.

To analyze structural changes that may occur during transcription, we extracted specific structures chosen by the *select* method at each nascent length. We viewed the family of selected structures at each length as RNAbow diagrams (Aalberts and Jannen, 2013), which revealed specific structural changes across the SRP RNA folding trajectory that differ between out-of-equilibrium and equilibrium datasets (Figure 2B-H, Figure S2B-E, Figure S2G-J). When 23-25 nts are free to fold in the cotranscriptional SHAPE-Seq predictions, we detect the formation of a 5’ helix containing 3 or more base pairs which persists through most of the folding pathway (Figure 2B-F). Interestingly, these 5’ helices differ from a previously inferred 5’ helix consisting of positions 4-10 paired to 16-22 suggested by the original enzymatic probing experiments (Wong et al., 2007). Instead, we consistently predict a 5’ helix where positions 3-8 are paired to 20-25 that we term helix 1 (H1). H1 is present for a large portion of the folding pathway and, based on our reconstructed states, can start to rearrange into the native long helical structure at length 110-111 nt (Figure 2B-G, Figure S2B-D, Figure S2G-I, SI Movie 1).

The next highly persistent structure that forms is a helix created when nts 53-55 pair with 60-62 to form the apical stem-loop of the native structure (Figure 2C-G, Figure S2B-E,G-J). We note, however, that our reconstructions do not predict the formation of 4 noncanonical interactions that are present in the crystal structure of the *E. coli* SRP RNA: C49-A76, A50-C75, G51-G74, and G52-A73 (Batey et al., 2000). We attribute this to the reliance of R2D2’s sample-and-select method on RNAstructure’s *partition* and *stochastic* functions which are not able to sample structures that contain noncanonical interactions, although portions of the cotranscriptional SHAPE-Seq reactivity matrix in this region show elevated reactivities indicating this region also likely does not close on the 30s timescale of our experiment. Despite these differences, R2D2 does reconstruct most of the mature SRP RNA sequence structure by length 117 (Figure 2H, Figure S2E,J).

Prior to folding into the mature structure, the sample-and-select method also predicts 3’ stem loop structures at various transcript lengths. One 3’ stem loop structure is between nucleotides 72 to 90, which we denote early helix 3 (eH3), and the next is between nucleotides 87 to 105, which we denote helix 3 (H3). Both eH3 and H3 locally sequester bases that need to be freed from these structures to form pairs in the mature structure. H3 was previously found by comparative analysis of SRP RNA sequences from diverse bacterial species and their phylogenetic tree, suggesting it may be an evolutionary conserved transient structural feature of the SRP RNA (Zhu et al., 2013). The presence of H1 and H3 present a significant structural barrier to cotranscriptional folding in that both must be broken to form the mature extended helical fold. We note however that H3 and eH3 are not predicted in every selected structure indicating that additional folding pathways could be possible.

### Sample-and-select models differ from approaches that do not use experimental data

Based on the ΔG folding trajectory, R2D2’s sample-and-select chooses structures that are higher in free energy than the MFE with or without experimental data at almost all lengths (Figure 2A, Figure S2A,F, Figure S3A, SI Movie 2). This is due to the difference between the R2D2 and the MFE objective functions: minimize a distance model that compares structures to experimental data, or minimize a free energy model, respectively. The addition of experimental data to the MFE prediction provides a structure more consistent with the experimental data (Figure S3B,C). Other than MFE approaches, one of the most widely used is KineFold, which simulates cotranscriptional folding given only an input sequence and a desired transcription rate (Xayaphoummine et al., 2005). In a comparison between 100 repetitions of KineFold and R2D2, KineFold predicted different folding pathways, with key differences in predictions of transient helices such as H1 and H3, and the location of the rearrangement (Figure S4).

### *A single point mutation disrupts the cotranscriptional rearrangement of the* E. coli *SRP RNA sequence*

R2D2 predictions show structural variation within H1 across the folding pathway (Figure 2, Figure S2, Figure 3A), which we hypothesized is due to the G-U pair within the otherwise G-C rich H1. We therefore mutated the native sequence to U21C to change the G-U pair into a G-C pair, to increase the stability of H1 and make the rearrangement into the final extended helix structure less favorable (Figure 3B). SRP RNA U21C cotranscriptional SHAPE-Seq reactivities (Figure 3C) contain higher reactivities within the loop of H1 throughout the folding pathway, suggesting it does not consistently refold into the native-like structure. In contrast, the equilibrium refolding experiment on the SRP RNA U21C mutant contains a visual drop in H1 loop reactivities around length 110 nt characteristic of the rearrangement into the final extended structure. R2D2 analysis of the cotranscriptional SRP RNA U21C dataset predicts the presence of H1 at all lengths of the folding pathway through length 112 nt (Figure 3D-H). The lack of H1 predictions at lengths 113-117 nts (Figure S5) may indicate that rearrangement of H1 is possible given the experimental data. Alternatively, this could be due to limitations in the Boltzmann distribution-directed sampling methods used, which is naturally biased towards sampling lower free energy structures making it difficult for the algorithm to choose out-of-equilibrium structures especially as the RNA length grows. To investigate this, we added to the selection pool structures sampled from the previous six lengths and extended them with unpaired nucleotides. With these additional structures, we find that rearrangement is not highly predicted at lengths 113-117 nt (Figure S5A-E). Notably, these predictions also showed the presence of H3 at the 3’ end of the RNA after lengths 103-105 nt that is present in the native sequence models as well. We also ran this sampling procedure on the native SRP RNA sequence as a control and found lengths 115-117 nt (Figure S5F) are predominantly predicted as rearranged as expected. Application of the standard R2D2 sample-and-select procedure to the SRP U21C equilibrium refolded datasets showed the presence of H1, but recovered the rearrangement into the final extended helical structure after length 109 nt (Figure 3D-H). Taken together this data demonstrates that a single point mutation can disrupt the SRP RNA cotranscriptional folding pathway and kinetically trap RNA into non-native intermediate structures.

**Figure 3.**
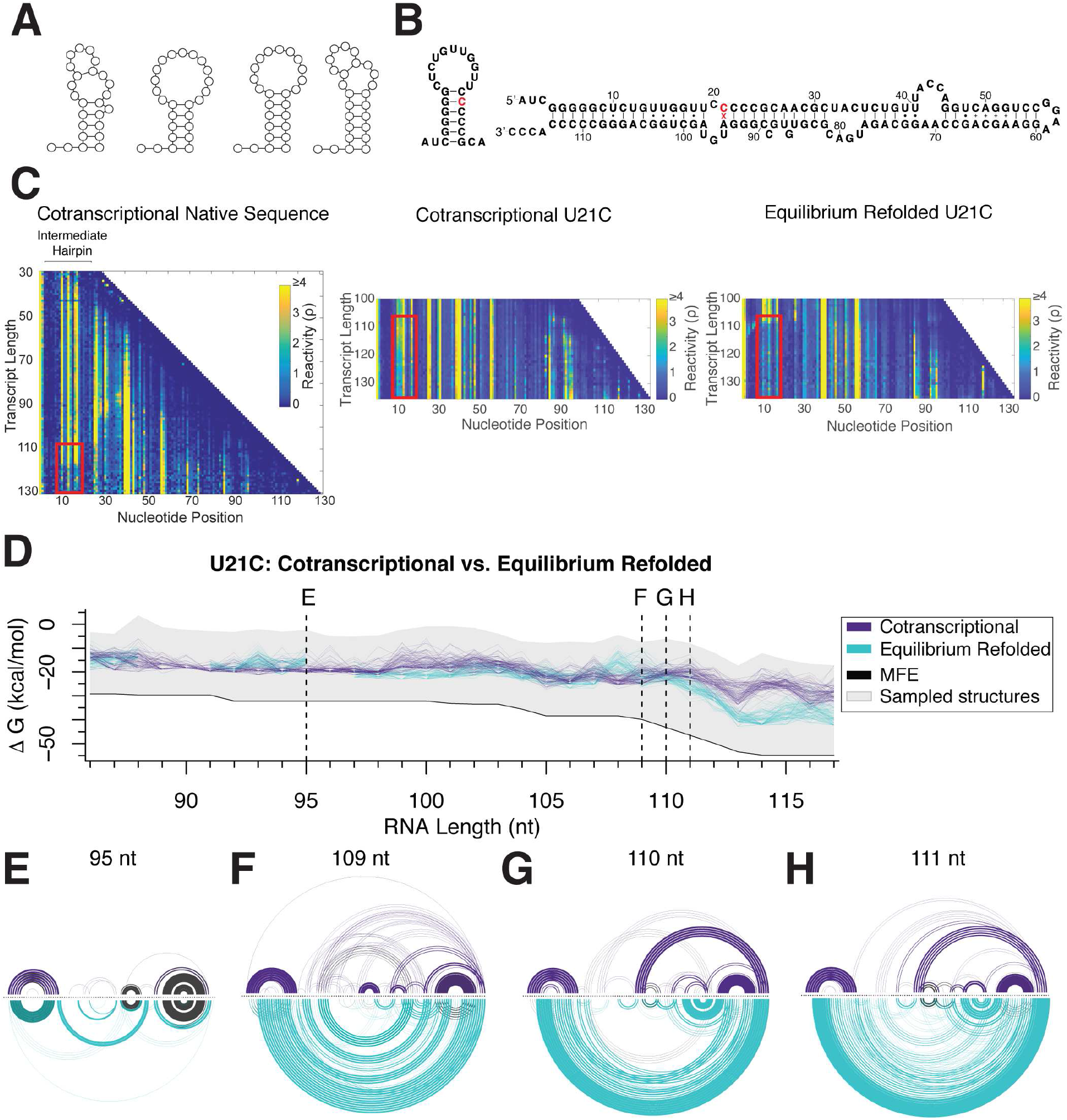
A single point mutation disrupts cotranscriptional rearrangement of the *E. coli* SRP RNA sequence. **(A)** Examples of 5’ helix (helix 1) variability in 2D predictions in the native sequence folding pathway indicate potential flexibility in the 5’ helix. **(B)** Diagram of the SRP RNA U21C mutation on the predicted 5’ helix and the secondary structure of the full length sequence. **(C)** Cotranscriptional SHAPE-seq reactivities from the native (left) sequence shows a drop in reactivities in nts 11-18 (red box) towards the end of the folding pathway indicating the rearrangement of helix 1 into the extended structure. The reactivity matrix for the SRP RNA U21C sequence (middle) has high reactivities in these positions throughout, indicating a disruption of the rearrangement with this point mutation. Equilibrium refolded SRP RNA U21C SHAPE-seq data (right) shows a drop in reactivities in this region, indicating that the disruption is only within a cotranscriptional folding regime. **(D)** Trajectory plot of R2D2 predictions for the U21C sequence following Figure 2. Structures over four lengths are highlighted: **(E)** 95 nt, **(F)** 109 nt, **(G)** 110 nt, and **(H)** 111 nt.

### *A single GU wobble is critical for the* E. coli *SRP RNA cotranscriptional rearrangement into the extended final fold*

Since the replacement of a single GU pair in the SRP RNA predicted H1 helix is enough to break the cotranscriptional rearrangement, we sought to test if reintroducing a GU pair in H1 would rescue the cotranscriptional rearrangement. We therefore designed a mutation (U21C, C22U, G93A) that reintroduces a GU wobble pair one position lower in the stem of H1 and maintains sequence complementarity between nt 22 and nt 93 (Figure 4A). Visually, the cotranscriptional SHAPE-Seq reactivity matrix for this mutant shows a drop in reactivities around length 118 nt, indicating that the nativelike rearrangement can occur cotranscriptionally (Figure 4B). In addition, when applied to this dataset, R2D2’s sample-and-select predicts that mutant model follows a similar folding pathway as the native sequence (Figure 4C,D), and that the rearrangement occurs at length 111 nts (Figure 4E). Overall this data point to the critical nature of the GU pair within H1 to facilitate the cotranscriptional rearrangement into the final extended helix structure.

**Figure 4.**
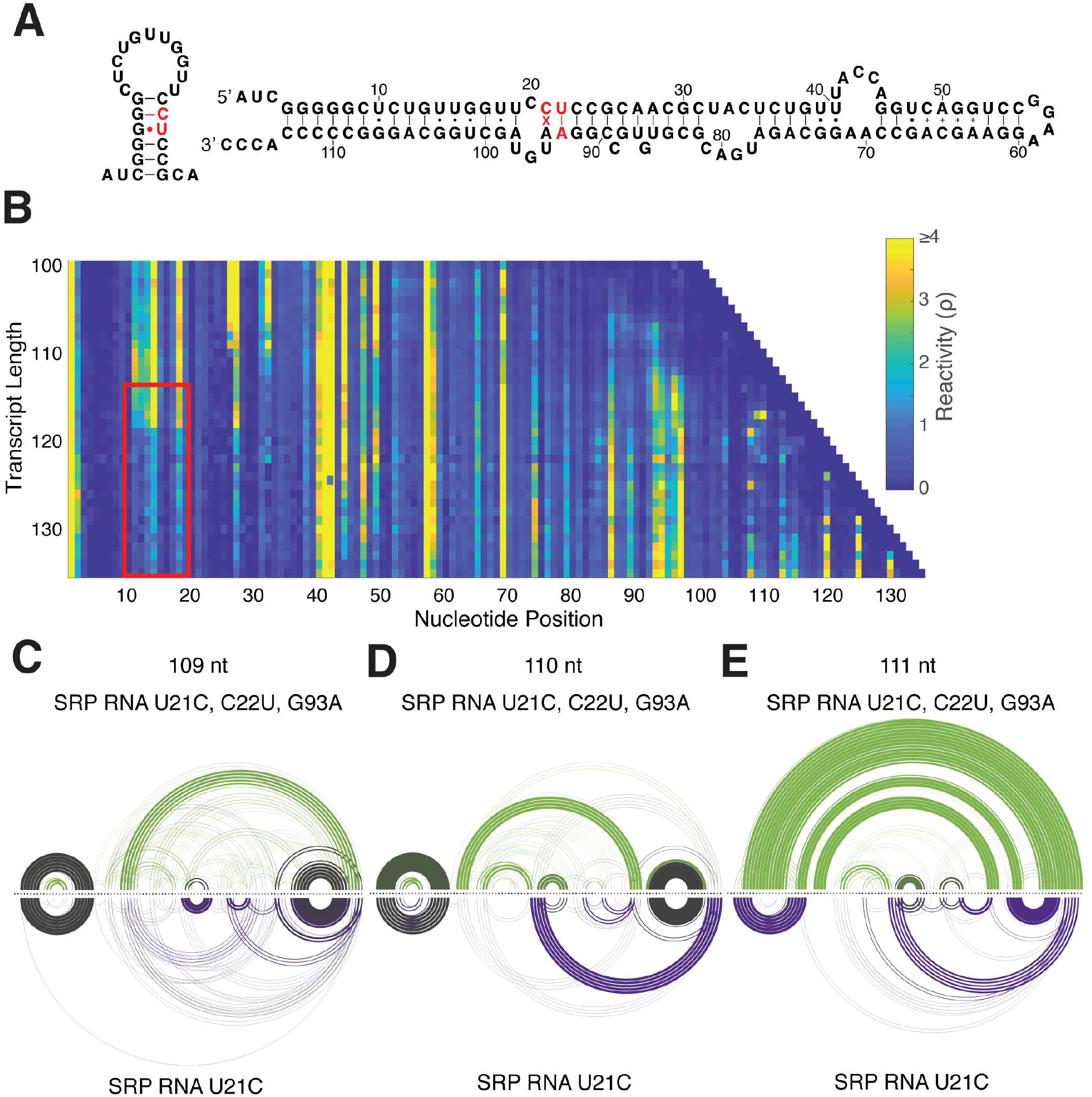
A rescue mutant of SRP RNA U21C confirms the importance of flexibility in helix 1. **(A)** Diagram of the rescue mutant U21C, C22U, G93A. The rescue mutant introduces a G-U pair in the middle of the SRP RNA U21C predicted H1 structure, overlaid on the predicted 5’ helix and extended full length structure. **(B)** Cotranscriptional SHAPE-seq reactivities from SRP RNA U21C, C22U, G93A. Red box indicates reactivities of some nucleotides spanning 11-18nt in helix 1 loop drop indicating cotranscriptional rearrangement into the extended structure. **(C-E)** RNAbow plots of SRP RNA U21C, C22U, G93A (green and top) and U21C (purple and bottom) R2D2 predictions following Figure 2. Three lengths are highlighted: **(C)** 109 nt, **(D)** 110 nt, **(E)** and 111 nt.

### Uncovering potential mechanisms of the SRP RNA cotranscriptional structural rearrangement with all-atom simulations

We next sought to determine the mechanism by which the SRP RNA cotranscriptionally rearranges and therefore the route by which the H1 GU pair facilitates this process. Paradoxically, H3 would be expected to impede the rearrangement process, as both it and H1 need to somehow unzip and hybridize together to form the native extended helix structure. We therefore focused on mechanisms by which the three hairpin consensus structure at 109 nt of cotranscriptional SHAPE-seq Replicate 1 (Figure 2F) can rearrange into the extended helix structure at 110 nt (Figure 2G). Four distinct potential transition pathways were identified: the inside-out (Figure 5A), kissing loop (Figure 5B), late toehold pathways (Figure 5C), and early toehold pathways (Figure 5D). We used all-atom molecular dynamics simulations to characterize the relative feasibility of each of the four proposed transition pathways from the stable folding intermediate containing H1 and H3 (Figure 6A) to the mature fold (Figure 6B). Each pathway suggests that the rearrangement mechanism initiate with a different set of specific base interactions (Figure 5). Therefore, to implement each pathway test, weak biasing forces were sequentially added in a specific order, starting at the initial proposed interaction, for each path to facilitate transitioning to the mature fold, with eight replicate simulations per path (Methods).

**Figure 5.**
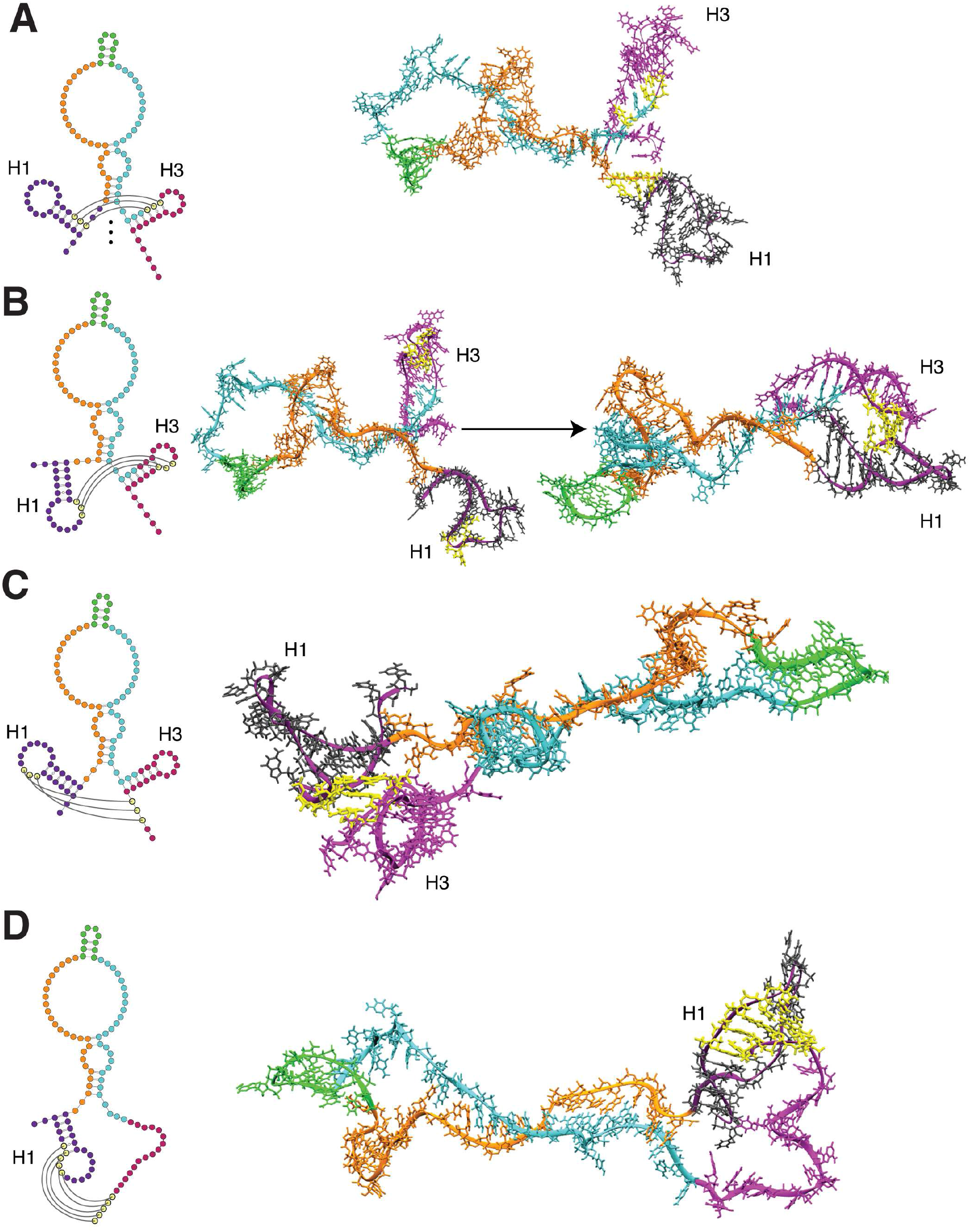
Snapshots of possible rearrangement mechanisms tested by 3D all atom simulations. R2D2 predicted secondary structures (left), were used as starting points for all atom MD simulations (right). For each rearrangement hypothesis, the pairing interactions that could seed the rearrangement into the native extended hairpin are indicated in yellow. Other bases are colored for convenient visualization: nts 1-25 (dark purple), 26-52 (orange), 53-62 (green), 63-96 (turquoise), and 97-117 (magenta). **(A)** The inside-out hypothesis whereby H1 and H3 progressively open and convert into the extended base pair configuration. **(B)** The kissing loop hypothesis where the loops of H1 and H3 begin the rearrangement process. **(C)** The late toehold hypothesis where bases 106-108 at the 3’ end of H3 seed the rearrangement through a toehold (yellow) with unpaired loop bases 9-11 of the 5’ hairpin. **(D)** The early toehold hypothesis where bases 106-110 seed the rearrangement through a toehold (yellow) with unpaired loop bases 7-11 of H1. The early and late toehold hypotheses differ in the structural state of the 3’ end before the rearrangement, with the late toehold hypothesis considering the rearrangement of the H3 structure in this region.

**Figure 6.**
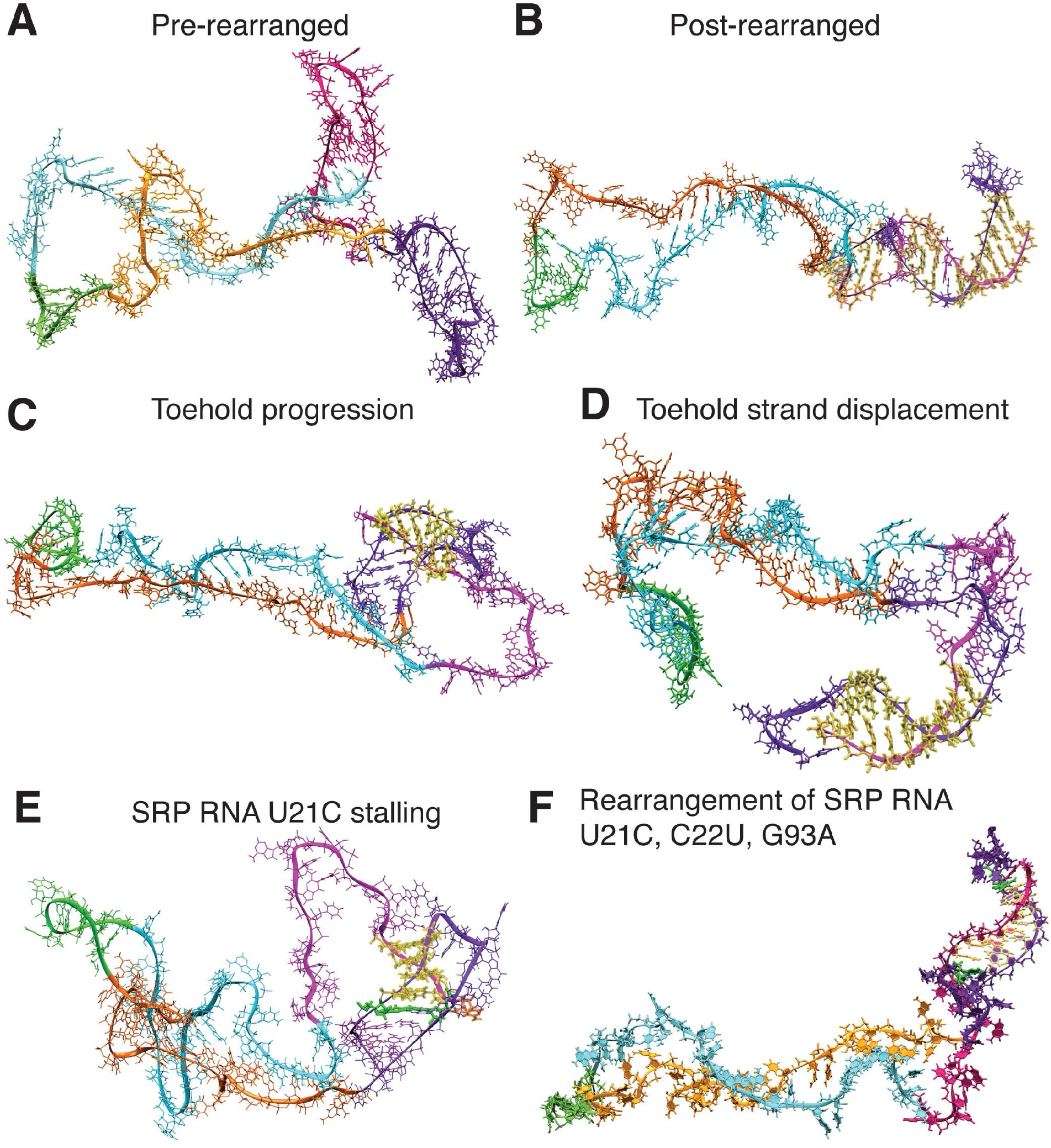
Snapshots of the toehold-mediated rearrangement pathway from molecular dynamics simulations constrained by R2D2 secondary structures. **(A)** Pre-rearranged structure with helix 1 (purple) and helix 3 (magenta) present. **(B)** Post-rearranged structure with new native base pairs (yellow) forming the extended helix. **(C)** Toehold progression to 6 bp of the native helix (yellow) requires unfolding of the weak 3’ hairpin (magenta). **(D)** Further elongation to a 9bp native helix (yellow) requires unfolding of the 5’ hairpin. **(E)** In the SRP RNA U21C mutant, the 5’ hairpin is stabilized by a GC basepair instead of GU (green). Even if a toehold is made to form (yellow), folding stalls as the G7-U21 basepair cannot be invaded even when modest biasing forces are applied. **(F)** In the SRP RNA U21C, C22U, G93A mutant, the rearrangement can occur and the GC basepair (green) can break.

The inside-out hypothesis involves simultaneously breaking H1 and H3 at their stems such that newly formed base-pairs form the native helix from the middle radiating outwards (Figure 5A, SI Movie 3). While it was technically possible to observe the inside-out pathway in the simulation, it would be extremely kinetically unfavorable since it would involve breaking two base-pairs for every one base-pair formed for a significant portion of the pathway (Figure S6, SI Movie 3). When more modest restraints were used, we found that this pathway could not spontaneously proceed. This indicated that this pathway was unlikely to be physiologically relevant and it provided an upper limit for how strong the applied restraints should be for all of the other transition pathways simulated.

The kissing loop pathway assumed that bases 17, 18 and 19 of the H1 loop and bases 98, 99, and 100 of the H3 loop form initial base pairs that seed the rearrangement (Figure 5B, SI Movie 4). These nucleotides were chosen because the resulting CG/GU/AU basepairs would produce a significantly stronger kissing-complex than ones composed of only GU/AU basepairs, analogous to the 2 bp kissing complex that drives MMLV genome dimerization (Li et al., 2006). The kissing loop was not able to form in all simulations of this pathway, even when each simulation was extended multiple times for 100 ns and the strength of the long-range restraints were doubled (Figure S6, SI Movie 4). This is because the mismatch in length of the two helical segments effectively prevents the bases from forming hydrogen bonds in the pretransition secondary structure, as they are geometrically constrained from aligning their hydrogen bond donors and acceptors.

Finally, the late toehold pathway assumes that bases 106-108, which are predicted to be in an unpaired strand at the 3’ tail of the RNA at the base of H3, initially pair with bases 9-11 in the H1 loop to forming a “toehold” interaction (Figure 5C, SI Movie 5). The initial toehold contacts were found to reliably form in 6/8 attempts as the 3’ tail of the nascent RNA is flexible and long enough to reach the loop of H1 (Figure S6). All simulations that formed the initial toehold contacts proceeded through the refolding pathway to the 110 nt structure.

An advantageous feature of the toehold mechanism is the favorability of the strand exchange process that proceeds in a break-one-form-one manner instead of break-some-form-one. Once identified as a plausible mechanism, we realized that this toehold-mediated strand-displacement can also be initiated earlier in the folding trajectory before H3 forms (Figure 5D). Simulations of the “early toehold” indicate that the absence of H3 actually speeds up the rearrangement due to the greater flexibility of the longer single-stranded 3’ tail, the lack of an energetic barrier posed by H3 (Figure 5D), and the increased number of bases available to form the initial toehold. Thus, the toehold-mediated strand-displacement mechanisms are much more plausible than the other pathways considered.

A detailed examination of the productive toehold-mediated folding pathways reveals several key architectural features that facilitate the rapid folding transition (SI Movie 5). Extension of the initial toehold seeding interaction through the full rearrangement requires fluctuations in the top of H1’s 11nt loop and into the stem (Figure 6C). H3, which is weaker than the GC rich H1 readily unfolds in the simulations after the first few base-pairs are formed, and the resulting increased single-strandedness further facilitates flexibility in hybridization with the H1 loop (Figure 6D). In addition, the formation of C7-G110 and C8-G109 base-pairs requires disruption at the top of H1’s stem, which is facilitated by partial opening of the GU base-pair. Only after C7-G110 and C8-G109 are formed is the H1 hairpin weak enough to open up, allowing the remaining basepairs of the native helix to align and zip-up in an energetically downhill process to form the fully extended fold (Figure 6B).

These results suggested that the SRP RNA U21C mutant removes the ability for H1 to fluctuate which is the root cause of the inability of this mutant to efficiently rearrange during transcription. To directly test this hypothesis, we also performed simulations of the SRP RNA U21C mutant, which revealed that the toehold can still form between bases 7-110 and 8-109 when restraints were applied, but the mutant cannot transition into the final folded state because of the increased stability of H1 (Figure 6E, Figure S6). The folding transition still stalled even when double-strength restraints were applied as it was still not enough to disrupt the C21-G7 basepair. Finally, simulations of the rescue mutant (U21C, C22U, G93A) confirm that restoring flexibility in the upper stem of H1 recovers the ability to transition to the mature fold, albeit at a slower rate due to the extra basepair that needs to be displaced to unfold H1 (Figure 6F, Figure S6).

Overall our all-atom MD simulations strongly suggest a toehold mediated strand displacement mechanism for refolding H1 into the final extended native helix and the critical importance of flexibility within the stem of H1 to enable toehold elongation into the fully rearranged extended helix.

## Discussion

We developed R2D2 to reconstruct nascent RNA folding at high resolution. Using the *E. coli* SRP RNA sequence as a model for studying cotranscriptional rearrangements, we show that the R2D2 sample-and-select approach both corroborates previous evidence for SRP RNA folding intermediates (Wong et al., 2007) and identifies additional secondary structural features that mediate folding. We then use these data driven secondary structure inferences as a starting point for all-atom MD simulations to test mechanistic hypotheses about how the RNA dynamically traverses intermediate states during transcription. While our *in vitro* conclusions directly apply to a truncation of the natively transcribed SRP sequence studied previously using enzymatic probing techniques (Wong et al., 2007), they could have potential implications for understanding the folding of cellular RNAs that must traverse through similar non-native structures during transcription to generate functional RNA structures.

The R2D2 approach builds off of elements from previously developed RNA folding algorithms, but is distinct in its methods towards the goal of reconstructing out of equilibrium folded states along a cotranscriptional folding pathway. Thus, R2D2 is naturally distinct from MFE prediction methods, which would not uncover the importance of H1 flexibility because of its stability in the SHAPE-directed MFE folding pathway (Figure S3, SI Movie 2). R2D2 uses a sample-and-select strategy to incorporate structure probing data into RNA folding models, which was inspired by a similar approach used in SeqFold (Ouyang et al., 2013). However, SeqFold uses a coarsegrained distance metric to predict a single structure that represents the data, while our distance metric uses the richness of the SHAPE-Seq reactivity scale to ultimately predict multiple structures that are all consistent with the data. In this way, the secondary structure aspect of R2D2 is similar to the recent development of the SLEQ (Li and Aviran, 2018) and Rsample methods (Spasic et al., 2017), although these methods are able to additionally estimate population levels of certain RNA structures which is not currently implemented in the R2D2 approach. Finally, R2D2 is distinct from algorithms that explicitly aim to predict cotranscriptional folding pathways from sequence alone such as KineFold (Xayaphoummine et al., 2005).

While powerful, R2D2 is not without limitations, many of which are inherent in the underlying algorithms used to sample possible structures. Specifically, there are currently no efficient methods to sample RNA structures with pseudoknots, non-canonical base pairs, or RNA-ligand/protein/RNA interactions (Ding et al., 2004; Tan et al., 2017). Structures that can be efficiently sampled are biased to the equilibrium Boltzmann distribution, which we try to overcome by sampling 150,000 states at each RNA length instead of the more commonly used 1,000-10,000 (Ding et al., 2004; Kutchko et al., 2015; Li and Aviran, 2018; Ouyang et al., 2013; Tan et al., 2017) (Figure S1). Since all-atom molecular dynamics was used to connect structural states uncovered by the secondary structure reconstruction method, it is inefficient to connect all possible sets of states together to reconstruct a full dynamic cotranscriptional folding pathway. However, our emphasis on using all-atom simulations to connect states is a unique feature of our approach to uncovering mechanisms of specific cotranscriptional rearrangements. Lastly, the predictions of R2D2 are only as good as the experimental data used. For example, the similarity between the cotranscriptional and equilibrium refolded folding pathway predictions for early lengths of the pathway in Figure 2 is expected, as the 30s duration of the cotranscriptional SHAPE-Seq experiment likely provides enough time for shorter length RNAs to equilibrate (Watters et al., 2016a). In addition, we have noticed that we typically observe lower reactivities towards the 3’ end of transcript lengths that could influence the computational inference of structures. Despite these limitations, we successfully generated a high resolution mechanistic model of the cotranscriptional folding of the *E. coli* SRP RNA sequence.

Our analysis of the SRP RNA cotranscriptional folding pathway suggests the formation of a structure containing non-native helices prior to the molecule rearranging into its native extended helical state. We focused these studies on a particular nonnative structure consisting of three helix elements in a row that was present in consensus R2D2 predictions: H1, the upper portion of the native helix, and H3 (Figure 2F, Figure 5A, Figure 6A). H3 may in fact be an evolutionary conserved element of the SRP RNA folding pathway, as it was previously identified using TRANSAT which incorporates phylogenetic sequence information (Zhu et al., 2013). Intriguingly, formation of H1 and H3 as cotranscriptional folding intermediates presents a conundrum for the folding pathway to adopt the native extended structure because these two helices sequester sequence elements that must somehow unwind and pair together. Using this intermediate structure as a starting point, we assessed four rearrangement mechanisms using all-atom MD simulations. Based on this analysis, we propose that the three-helix structure can efficiently rearrange into the single extended helix through a toehold mediated strand exchange mechanism (Seelig et al., 2006; Yurke et al., 2000; Zhang et al., 2007). In this mechanism, the single stranded nucleotides 3’ of H3 serve initially as a ‘toehold’ and bind to a complementary portion in the H1 loop and seed the rearrangement. Once the toehold is seeded, H1 and H3 can then exchange base pairs in a break-one-form-one pathway that avoids large energetic barriers and thus can efficiently rearrange to the mature structure. Our simulations of the toehold strand displacement mechanism also highlights the importance of conformational flexibility in these intermediate helices, especially H1, for efficient toehold-mediated strand displacement. The importance of H1 flexibility is supported by the observation that the single point mutation of the SRP RNA, U21C, was enough to experimentally kinetically trap the rearrangement by stabilizing H1, which could then be rescued by reintroducing this flexibility in H1 by two additional point mutations.

While the toehold mediated strand exchange mechanism is compelling and what we focused on in this study, we note that alternative folding pathways are possible. Even within toehold-mediated mechanisms, multiple toehold initiation points and rearrangement starting from eH3 or other 3’ structures instead of H3 are possible. The entire loop of H1 is base paired within the native fold, indicating that there are many positions that could serve as an initial toehold nucleation point. The large size of the H1 loop could also be important for the increased flexibility of these bases for toehold nucleation as well as exposing a large sequence target to capture the many alternate transient 3’ end structures to seed the rearrangement process.

Overall, it could be that the SRP RNA sequence has evolved to allow multiple toehold strand exchange mechanisms in order to prevent the kinetic folding trap imposed by H1. Recently it has been shown that many natural RNAs contain long-range interactions in the cell, some of which occur over 1 kb away (Lu et al., 2016). Given the propensity of RNAs to form local structures cotranscriptionally, toehold-mediated strand displacement could be one of the most efficient ways for RNA molecules to undergo large scale rearrangements. Detailed studies of toehold strand displacement reactions in designed *in vitro* systems have demonstrated their speed, with reactions proceeding with rates on the order of 10^6^/M/s for a bimolecular strand exchange reaction (Šulc et al., 2015; Zhang and Winfree, 2009). In addition, the elementary steps of strand exchange can be inferred to occur on the μs timescale, which is orders of magnitude faster than the ms timescales of nucleotide incorporation during transcription (Roberts et al., 2008). Intriguingly, to the best of our knowledge, the observation of this mechanism within the *E. coli* SRP RNA cotranscriptional folding pathway could be the first observation of toehold mediated strand displacement in a naturally-derived RNA sequence, and suggests this may be an important mechanism yet to be broadly uncovered in biology.

While this manuscript was being prepared, an independent study of the cotranscriptional folding pathway of the same *E. coli* SRP sequence was performed using single-molecule optical tweezers techniques (Fukuda et al., Submitted). A close inspection of the data from this study revealed several structural features consistent with our findings including the formation of H1, the presence of structures that could form near the 3’ end transcription of the molecule prior to rearrangement which matches our prediction of H3 (e.g. denoted as H4 in that study), and an independent observation that the SRP RNA U21C mutation introduces a kinetic trap at the single molecule level (i.e. numbered as U18C in that study). Furthermore, the datasets from Fukuda *et al*. also display a fascinating occurrence of ‘hopping dynamics’ near the structural/folding rearrangement transition which consist of large scale fluctuations in RNA end-to-end distances. These hopping dynamics could be originated from the molecular search for toehold seeding interactions, or strand exchange attempts that open structural elements before the rearrangement. The overall agreement of these two complementary studies highlights the power of combining them to give a deeper and more completed mechanistic view of cotranscriptional RNA folding.

These studies begin to address a fascinating question for cellular biology – how do RNAs efficiently fold into functional states and exit the kinetic traps imposed by the polarity and timescale of cotranscriptional folding in the cell? While we cannot rule out the possibility of other interactions and cellular processes facilitating these rearrangements, it is possible that these molecular sequences utilize some of the same principles of the rearrangement pathways studied here to find these structures. Overall these studies are starting to uncover deeper principles of RNA cotranscriptional folding that may still be hidden, but are critically important, and occur every time an RNA is transcribed.

## Acknowledgements

We thank Eric Siggia (Rockefeller University) for conversations that laid the foundations for this work. We thank Dave Matthews (University of Rochester) for many helpful conversations on RNA structure prediction. We thank Daniel Aalberts (Williams College) for assistance with plotting RNAbow plots for figures and movies, and Katherine Berman for helping to clone SRP RNA sequence mutants. We also thank S. Yan, S. Fukuda, and C. Bustamante (University of California, Berkeley) for helpful discussions. Support for this work was provided by the Tri-Institutional Training Program in Computational Biology and Medicine (via NIH training grant T32GM083937 to AMY), through the National Institute of General Medical Sciences of the National Institutes of Health by a New Innovator Award [grant number 1DP2GM110838 to JBL], the National Science Foundation [grant MCB1651877 to AAC], and Searle Funds at The Chicago Community Trust [to JBL]. This work used the Extreme Science and Engineering Discovery Environment (XSEDE) [allocation TG-MCB140273 to AAC], which is supported by National Science Foundation grant number ACI-1548562. This research was supported in part through the computational resources and staff contributions provided for the Quest high performance computing facility at Northwestern University which is jointly supported by the Office of the Provost, the Office for Research, and Northwestern University Information Technology. The content is solely the responsibility of the authors and does not necessarily represent the official views of the National Institutes of Health.

## Methods

### Contact for Reagent and Resource Sharing

Further information and requests for resources and reagents should be directed to and will be fulfilled by the Lead Contacts, Julius Lucks and Alan Chen.

### Method Details

#### Cotranscriptional and Equilibrium-Refolded SHAPE-Seq

The SRP RNA sequence used to generate mutants was previously described (Watters et al., 2016a; Wong et al., 2007) and substitutes a 24 nt leader sequence with AUC at the 5’ end (RNA Mapping Database accession codes: SRPECLI_BZCN_0001, SRPECLI_BZCN_0002, SRPECLI_BZCN_0003, SRPECLI_BZCN_0004). DNA templates for cotranscriptional SHAPE-Seq were prepared as previously described (Watters et al., 2016a). DNA templates specifically targeted transcript lengths 101 to 136. Cotranscriptional SHAPE-Seq experiments were performed as previously described, except that EcoRI_E111Q_ was included at 800 nM during *in vitro* transcription instead of 500 nM (Watters et al., 2016a).

#### Reactivity Calculation

Quantification of reactivities from cotranscriptional SHAPE-Seq data was done using Spats v.1.0.1 (http://luckslab.github.io/spats/) as previously published (Watters et al., 2016b). The θ reactivities outputted by Spats were converted to *ρ* reactivities to allow for direct comparison of SHAPE probe accessibility between intermediate lengths of RNAs (Watters et al., 2016a). *ρ* is equivalent to *θ* multiplied by the length of the RNA and the average of *ρ*’s in one RNA is equal to 1 (Watters et al., 2016b). For cotranscriptional predictions where RNA polymerase occludes the last ~14 nts from folding (Komissarova and Kashlev, 1998; Watters et al., 2016a), ρ reactivities were trimmed by 14 nts and renormalized such that the reactivities average to 1. This trimming was not done for equilibrium-refolded predictions because there should be no occlusion from RNA polymerase.

#### R2D2’s sample-and-select for reconstructing secondary structures from experimental data

The R2D2 sample-and-select method was first developed to predict the equilibrium fold of a single RNA using equilibrium SHAPE-Seq data. A crucial step was to establish a method to *select* structures that are most consistent with the experiment. SHAPE-Seq reactivities, ρ, are values ≥ 0 that reflect the structural state of each nucleotide: ρ = 0 corresponds to a nucleotide that is present in a structured context (such as a base pair or stacking interaction), while ρ > 1 represents a nucleotide that is present in a flexible context (such as an unpaired region) (Bindewald et al., 2011). Thus ρ values most naturally correspond to a representation of the un-paired state of each nucleotide in an RNA secondary structure, which can be represented by a binary vector (*u* for ‘un-paired’) containing 0 if a nucleotide is paired and 1 if a nucleotide is un-paired (Figure 1C). Comparison between the ρ vector of reactivity data at a specific transcript length, and the *u* vector for a specific structure that is possible to occur at that length can then be easily made with a metric that reflects their distance from each other (Figure 1C, Table S1).

We developed and tested six functions to calculate the distance between a SHAPE-Seq reactivity spectra and a given RNA secondary structure (Figure 1C, Table S1). Each distance function is of the form

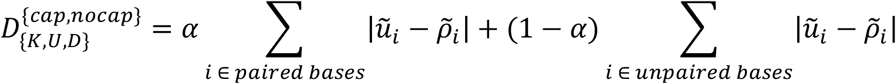

where 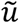 is a vector calculated from the *u*-vector of a specific RNA secondary structure, and 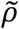 is calculated from the experimental SHAPE-Seq reactivity data ρ vector. Reactivity is inherently a measure of openness of an RNA structure, and low reactivity may not be due only to base pairing, but can be caused by other structural constraints such as stacking (Bindewald et al., 2011). To account for this, we incorporated a weighting between single-stranded and paired bases in sampled structures, *α*, which is used to adjust the contribution to the distance from positions that are predicted to be paired.

Since unpaired vectors and *ρ* vectors are different types of data (binary vs. continuous) and on different numerical scales, we explored three different ways to calculate their differences specified by K, U and D, which specify the way 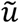 is calculated:

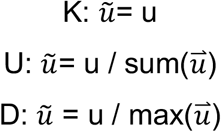

Since certain RNA folds can result in *ρ* values that are much larger than one (McGinnis et al., 2012), we also explored ways to cutoff *ρ* values at a maximum value. This is specified by the indices *cap* or *nocap* which determine the way 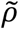 is calculated with *cap* meaning *ρ* is capped at a *ρ_max_* value 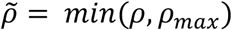, and *nocap* meaning the full *ρ* value is used.

The distance metrics above can be used to *select* structures from a candidate set that are most consistent with the observed experimental reactivity data by choosing the minimum distance structure(s) at every length (Figure 1C). To generate a candidate set of structures, the *sample* method statistically samples structures with a large sample size using the *partition* and *stochastic* functions of the RNAstructure suite of computational secondary structure prediction tools (Reuter and Mathews, 2010). The *partition* method takes as an input the RNA sequence and folding parameters, and uses them to calculate the secondary structure partition function for that sequence. The partition function for an RNA sequence describes how a population of RNA molecules with that sequence partitions into an ensemble of different structures in equilibrium, with each structure occurring with a Boltzmann probability. The *stochastic* method then uses this partition function to stochastically generate RNA structures according to their equilibrium Boltzmann probabilities – i.e. lower free energy structures are generated more frequently than higher free energy structures. Thus repeated application of the *stochastic* method can generate a set of possible candidate structures the RNA molecule may exist in during the experiment.

The goal of the *sample* method is to generate the greatest amount of structural diversity possible to allow more choices for the *select* method. An initial test of the degree to which the stochastic method can generate novel structures revealed that the method did not converge on exhausting the possibilities of different RNA structures even after 150,000 structures were drawn (Figure S1). This is not surprising since the free energy landscapes of RNA secondary structures are known to have a shallow density of states near the minimum free energy structure (Chen and Dill, 2000) indicating there are many possible RNA structures that are low in free energy and would be sampled frequently by the *stochastic* method. To circumvent this and still generate a diverse array of candidate structures without the computational burden of generating millions of structures, we used two additional variations of the sampling procedure that used experimental SHAPE restraints to calculate a modified partition function from which we could sample. The first, called SHAPE-directed sampling, used the *partition* method’s ability to incorporate SHAPE reactivities as effective free energy terms in the partition function calculation (Kutchko et al., 2015; Reuter and Mathews, 2010). The second, called SHAPE-forced sampling, used a SHAPE reactivity cutoff, *ρ_c_*, to force nucleotides with reactivities greater than this value to be single-stranded in the partition function calculation. In total, the *sample* method consisted of sampling 50,000 structures from each of these methods for a total of 150,000 structures which acted as the candidate set for the *select* method. We note that even though the *sample* method uses SHAPE reactivity data to generate part of the candidate set, these are not guaranteed to be chosen as most consistent with the data by the *select* method. Rather, they are included to increase the diversity of the candidate set.

Software implementing this method were run with Python 2.7.12 through Anaconda 2.4.1 (64-bit) and R version 3.2.2. Images and movies were made with ffmpeg version 3.1.3, ImageMagick 7.0.3-0 Q16 x86_64, and iMovie v10.1.2. Version 5.8.1 of RNAstructure was used for the *partition* and *stochastic* methods, and VARNA version 3.9 was used to visualize RNA secondary structures. See “Data and Software Availability” for location of code used in this study.

#### Benchmarking

Best parameter values were determined through a grid search of 10,404 parameter sets: all combinations of 0.7 to 4.1 by 0.1 for *ρ_c_*, 0.7 to 4.1 by 0.1 for *ρ_max_*, and 0 to 1 by 0.1 for *α*. The best parameter set(s) was determined as the parameter set(s) with the largest sum of *F*_1_ scores (F-scores) for 18 previously published equilibrium-refolded SHAPE-Seq datasets on 6 RNAs of known crystal structures (three replicates) and no pseudoknots since RNAstructure cannot sample structures with pseudoknots (Loughrey et al., 2014). F-score is defined as follows:

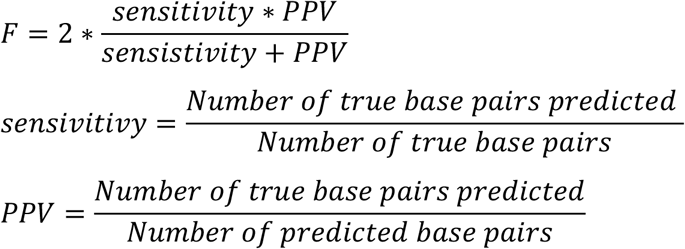

For every parameter set, we sampled 50,000 structures for each of the three sampling methods, for a total of 150,000 structures (see *“R2D2’s sample-and-select for reconstructing secondary structures from experimental* data”). For each benchmark RNA and dataset, the minimum distance structure was calculated and F-score determined from the prediction and the known structure. The sum of F-scores across the panel of RNAs and datasets was then reported for that parameter set. If multiple minimum distance structures were found, then the average of their sum of F-scores were used to find the best parameter set. We ran the benchmarking for each of the 6 distance equations. *ρ_max_* is not used when no reactivity capping is used, so only 306 parameter sets were tested in these cases.

We found two different metrics were the best performing across all distance functions: *D_K,cap_* with *ρ_c_* = 3.5, *ρ_max_* = 1.0 *or* 0.9, and *α* = 0.8 as well as *D_D,cap_* with *ρ_c_* = 3.5, *ρ_max_* = 1.0, and *α* = 0.8. These two each had an average F-score of 86.32% for the 18 RNA datasets in the panel (Table S1). From this set, we chose as our parameter set *D_K,cap_* with □_*c*_ = 3.5, *ρ_max_* = 1.0, and *α* = 0.8, which gives a higher weight to paired positions in the sampled structures as expected, and matches common interpretations of ‘high’ reactivity values being greater than 1. We note that this is mathematically equivalent to *D_D,cap_*’s best parameter set.

We also compared the best results from the sample-and-select method to SHAPE-restrained secondary structure predictions using the same data on the same RNA panel using the *Fold* method of RNAstructure (Table S1). In aggregate, the sample-and-select method (average F-score of 86.32%) does not perform better than RNAstructure-Fold with SHAPE restraints (average F-score of 88.95%), but does perform better than RNAstructure-Fold without SHAPE restraints (average F-score of 77.51%). Interestingly R2D2’s sample-and-select method did outperform on the *E. coli* TPP riboswitch in terms of sensitivity, PPV, and F-score for all replicates (Table S1). While the accuracy of our sample-and-select method applied to equilibrium RNA structure prediction is not overall better than the best equilibrium structure prediction algorithms given the same data, it was designed to find RNA secondary structures consistent with structural probing data from out-of-equilibrium RNA folds and thus can be used to reconstruct a complete secondary structure cotranscriptional folding pathway of an RNA.

Software implementing this method were run with Python 2.7.11 through Anaconda 2.3.0 (64-bit). Version 5.6 beta of RNAstructure was used for the *partition* and *stochastic* methods, and VARNA version 3.9 was used to visualize RNA secondary structures. See “Data and Software Availability” for location of code used in this study.

#### Application to cotranscriptional SHAPE-Seq data

We applied the method described in “*R2D2*’s *sample-and-select for reconstructing secondary structures from experimental data*” to each length of cotranscriptional SHAPE-Seq data available with the parameter set found in “*Benchmarking*”. For each structure predicted, free energies were calculated using RNAstructure-efn2.

Software implementing this method were run with Python 2.7.12 through Anaconda 2.4.1 (64-bit) and R version 3.2.2. Images and movies were made with ffmpeg version 3.1.3, ImageMagick 7.0.3-0 Q16 x86_64, and iMovie v10.1.2. Version 5.8.1 of RNAstructure was used for the *partition, stochastic, efn2, and ct2dot* methods. RNAbows was used to visualize R2D2 2D predictions (Aalberts and Jannen, 2013). See “Data and Software Availability” for location of code used in this study.

#### Minimum free energy folding pathway prediction

Each length of the SRP RNA sequence was folded with RNAstructure-Fold method without SHAPE restraints to generate the minimum free energy folding pathway. Images of the minimum free energy structures were made into a movie with RNAstructure-draw and ffmpeg. Free energy calculations were done with RNAstructure-efn2. The SHAPE-directed MFE folding pathway prediction was done similarly, but with ρ reactivities and m = 1.1 and b = −0.3 for lengths where SHAPE data was available in specified datasets.

Software implementing this method were run with Python 2.7.12 through Anaconda 2.4.1 (64-bit). Images and movies were made with ffmpeg version 3.1.3, ImageMagick 7.0.3-0 Q16 x86_64, and iMovie v10.1.2. Version 5.8.1 of RNAstructure was used for the *Fold* method to predict MFE structures, and *draw* methods (used for SI Movie 2) for drawing structures. RNAstructure version 6.0’s StructureEditor executable was used to draw structure outlines in Figure S3B,C. See “Data and Software Availability” for location of code used in this study.

#### KineFold predictions

KineFold cotranscriptional folding pathway predictions were done using the KineFold executable with ‘co-transcriptional fold’ with a new base added every 20 ms, no pseudoknots, and freely crossing entanglements. KineFold executable was used and can be downloaded from: http://kinefold.curie.fr/download.html. For each structure in KineFold’s .rnm output, the free energy was calculated using RNAstructure-efn2. See “Data and Software Availability” for location of code used to run KineFold and analyze .rnm output.

#### Using R2D2 predictions to inform all-atom folding pathway simulations

In order to assess the feasibility of the different hypothetical folding pathways in the full three-dimensional context of the folded RNA, the R2D2 secondary structures were used to restrain all-atom molecular dynamics simulations of each proposed transition pathway. Base-pair constraints for the pre- and post-folding transition were defined using the consensus (base pairs that occur in ≥ 50% of the 100 iterations) R2D2 secondary structures at length 109 and 110 nt respectively. In order to avoid overconstraining the simulation, only those base-pairs that occurred in over 50% of the reconstructions were enforced with explicit folding restraints. It should be noted that non-restrained bases can still form base-pairs according to the all-atom energy potential. While all pathways start from the same 109 nt folding intermediate (Figure 2F, Figure 5A,B,C,D), each pathway then dictates a unique order in which the base pairing pattern must rearrange to arrive at the final 110 nt native fold (Figure 2G). All-atom simulations employed the GROMACS 2016 software package (Abraham et al., 2015), using the Amber-99 force field (Wang et al., 2000) with Chen-Garcia modifications for RNA bases (Chen and García, 2013), the Case et al modifications for the backbone phosphate (Steinbrecher et al., 2012), the TIP4P-EW water model (Horn et al., 2004), and the Joung & Cheatham parameters for potassium chloride ions (Joung and Cheatham, 2008).

Simulations employed truncated dodecahedral boxes of ~15 nM radius, containing the 110 base RNA, 74,428 TIP4P-EW H_2_O’s, 1,559 K^+^ and 1,450 Cl^-^ ions to mimic 1M excess salt conditions to give a total of 304,265 atoms. Long-range interactions beyond 10 Angstroms were calculated using PME with a grid size of 0.16 nm. A constant pressure of 1 atm was maintained using the Berendsen barostat (Berendsen et al., 1984) with a time constant of 1.0 ps, and a constant temperature of 450K was maintained using the V-rescale thermostat (Bussi et al., 2007) with a time constant of 0.1 ps. The leapfrog Verlet integrator with a 2 fs timestep was used, with the total production length of each simulation being 100-500 ns according to the scheme below, leading to a cumulative total of >5 μs of simulations.

Base-pairs were restrained using a piecewise flat-bottomed harmonic restraint of strength 0.5 kcal/mol between central H-bond donor/acceptor of natively paired bases. This restraint becomes linear at distances greater than 4 Angstroms. The strength and distance dependence of the restraints was chosen to be strong enough to facilitate formation of long-range interactions in ~100 ns simulations, but not strong enough to significantly unfold other sections of the RNA in the process. Elevated temperatures were used to increase RNA flexibility and reduce the amount of computational time needed to sample each proposed transition pathway. This arrangement ensured that individual folding attempts would simply stall if two restrained bases could not physically get close enough to form a new basepair in the 3D context of each folding intermediate.

The 110 nt RNA chain was initially equilibrated until all base pairs observed in >50% of the stable folding intermediate (109 nt R2D2 2D prediction) were stably formed. At this point, new restraints from the 110 nt natively folded transcript were added 2-3 base-pairs at a time. Each new set of restraints were simulated at least 10 ns until they were successfully formed, at which point the next set of new restraints were added. This cycle was repeated until all bases were successfully paired in the RNA’s native fold. Simulations that had still not achieved any new base-pairs within 5 successive cycles (i.e. 50 ns) after adding restraints were considered “stalled” and not simulated any further. Eight separate attempts were made to simulate each of the 4 proposed pathways and two mutants studied. Each individual folding trajectory therefore ranged from 100-500 ns depending on stalling, and successful folding pathways exhibited at least 6/8 successfully folded trajectories while pathways deemed “unfeasible” always exhibited zero successful attempts.

Four potential folding pathways were simulated. In the “inside-out” pathway, the formation of the extended native helix proceeds by extending the predicted central helix along its axis, unraveling H1 and H3 during this progression, and eliminating the need for forming an initial long-range contact between the RNA ends (Figure 5A). In the “kissing loop” mechanism, it was proposed that complimentary, unpaired loop bases within the H1 and H3 hairpins could form an initial long-range “kissing complex”, which could then seed formation of the hybrid helix in a strand rearrangement process (Figure 5B). This hypothesis is attractive because kissing-loop interactions are known to be rapid and stable ways to form long-range RNA interactions in RNA gene regulation and retroviral replication (Kolb et al., 2000; Paillart et al., 2004). A toehold strand exchange mechanism was also explored, in which the free 3’ end of the nascent RNA chain initially hybridizes with unpaired bases in the loop of H1, seeding a sequential unfolding pathway where strands of H1 and H3 are exchanged with each other to rehybridize into the final extended native helix (Figure 5C). Finally, we also explored the “early toehold” mechanism which could initiate at different exposed bases of H1 before H3 is fully formed (Figure 5D).

## Data and Software Availability

R2D2 will be freely available here: https://github.com/LucksLab/R2D2

